# Nitric oxide signaling controls collective contractions in a colonial choanoflagellate

**DOI:** 10.1101/2022.02.14.480389

**Authors:** Josean Reyes-Rivera, Yang Wu, Benjamin G. H. Guthrie, Michael A. Marletta, Nicole King, Thibaut Brunet

**Affiliations:** Howards Hughes Medical Institute and the Department of Molecular and Cell Biology, University of California, Berkeley, CA, USA; Department of Chemistry and the Department of Molecular and Cell Biology, University of California, Berkeley, Berkeley, CA, USA; Institut Pasteur, Université de Paris, Department of Cell Biology and Infection and the Department of Developmental and Stem Cell Biology, F-75015 Paris, France

## Abstract

Although signaling by the gaseous molecule nitric oxide (NO) regulates key physiological processes in animals, including contractility^1-3^, immunity^4,5^, development^6-9^ and locomotion^10,11^, the early evolution of animal NO signaling remains unclear. To reconstruct the role of NO in the animal stem lineage, we set out to study NO signaling in choanoflagellates, the closest living relatives of animals^12^. In animals, NO produced by the nitric oxide synthase (NOS) canonically signals through cGMP by activating soluble guanylate cyclases (sGCs)^13,14^. We surveyed the distribution of the NO signaling pathway components across the diversity of choanoflagellates and found three species that express NOS, sGCs, and downstream genes previously shown to be involved in the NO/cGMP pathway. One of these, *Choanoeca flexa*, forms multicellular sheets that undergo collective contractions controlled by cGMP^15^. We found that treatment with NO induces sustained contractions in *C. flexa* by signaling through an sGC/cGMP pathway. Biochemical assays show that NO directly binds *C. flexa* sGC1 and stimulates its cyclase activity. The NO/cGMP pathway acts independently from other inducers of *C. flexa* contraction, including mechanical stimuli and heat, but sGC activity is required for contractions induced by light-to-dark transitions. The output of NO signaling in *C. flexa* – contractions resulting in a switch from feeding to swimming – resembles the effect of NO in sponges^1-3^ and cnidarians^11,16,17^, where it interrupts feeding and activates contractility. These data provide insights into the biology of the first animals and the evolution of NO signaling.

## RESULTS

### *C. flexa* encodes both NOS and sGC

*C. flexa* is a colonial choanoflagellate that forms concave sheets capable of global inversion of their curvature through collective contractility^15^. *C. flexa* inversion mediates a trade-off between feeding and swimming: relaxed colonies (with their flagella pointing inside) are slow swimmers and efficient feeders, while contracted colonies (with their flagella pointing outside) are inefficient feeders but fast swimmers^15^ (Fig. 1A). Inversion has been shown to be induced by light-to-dark transitions through inactivation of a rhodopsin-phosphodiesterase (Rho-PDE) and accumulation of cGMP^15^ (Fig. 1B). The involvement of cGMP in collective contraction in *C. flexa* and the connection between NO/cGMP signaling and tissue contraction in early-branching animals^18^ led us to investigate whether NO signaling might exist and regulate sheet inversion in *C. flexa* (Fig. 1B).

**Figure 1.**
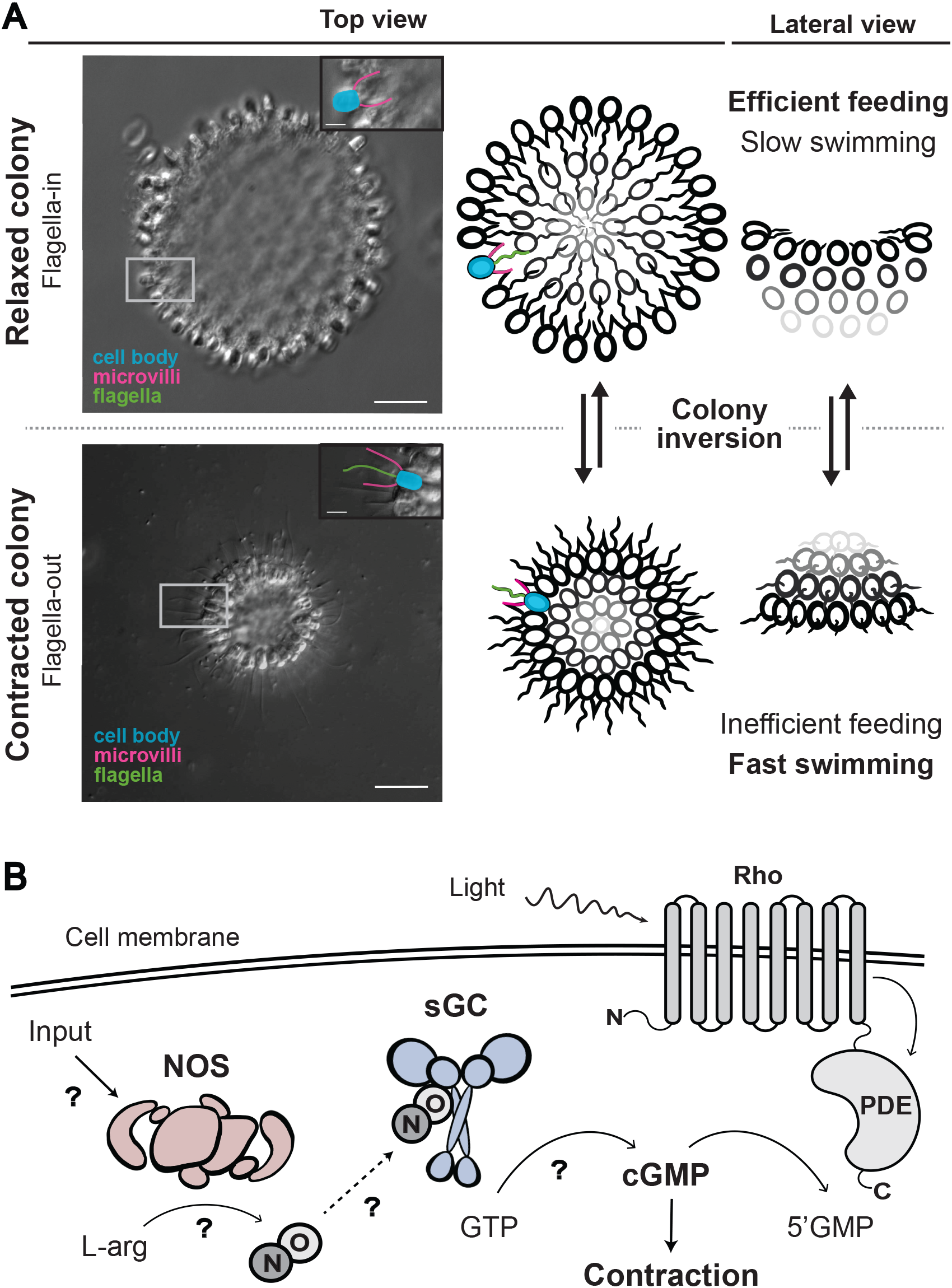
The inversion behavior of *C. flexa*, its control by light and hypothesized control by NO. **(A)** Micrograph (left) and graphical depiction (right) of *C. flexa* inversion behavior. Cells are linked by their collars and form a cup-shaped monolayer or sheet; scale bar: 15 μM. Insets: Pseudo-colors highlight the characteristic morphological features of choanoflagellates and the orientation of the flagella (flagella from relaxed colony not shown, out of focus); scale bar: 5 μM. Relaxed colonies (top half) have their flagella pointing towards the inside of the colony and are efficient feeders and slow swimmers. Contracted colonies (bottom half) have their flagella pointing towards the outside of the colony and are inefficient feeders but fast swimmers. **(B)** *C. flexa* colony inversion is controlled by light-to-dark transitions, mediated by a rhodopsin-phosphodiesterase fusion protein (Rho-PDE) upstream cGMP signaling. In the presence of light, Rho-PDE is active and constantly converting cGMP into 5’GMP. We hypothesized that NO/cGMP signaling might also be able to induce inversion. Mechanisms tested in this paper are highlighted with question marks: a primary input activates the nitric oxide synthase (NOS), which converts L-arginine into NO and L-citrulline. NO diffuses away and activates soluble guanylate cyclase (sGC) which convert GTP into cGMP, causing colony contraction.

By examining the genomes of two choanoflagellates (*M. brevicollis*^19^ and *S. rosetta*^20^) and the transcriptomes of *C. flexa*^15^ and 19 other choanoflagellates^21^, we found that five choanoflagellates encode NOS homologs and 10 encode sGC homologs (Fig. 2A). Of these, three species – *S. infusionum, C. perplexa*, and *C. flexa* – express both NOS and sGCs, suggesting that some aspect of the physiology of these organisms might be regulated by NO/cGMP signaling.

**Figure 2.**
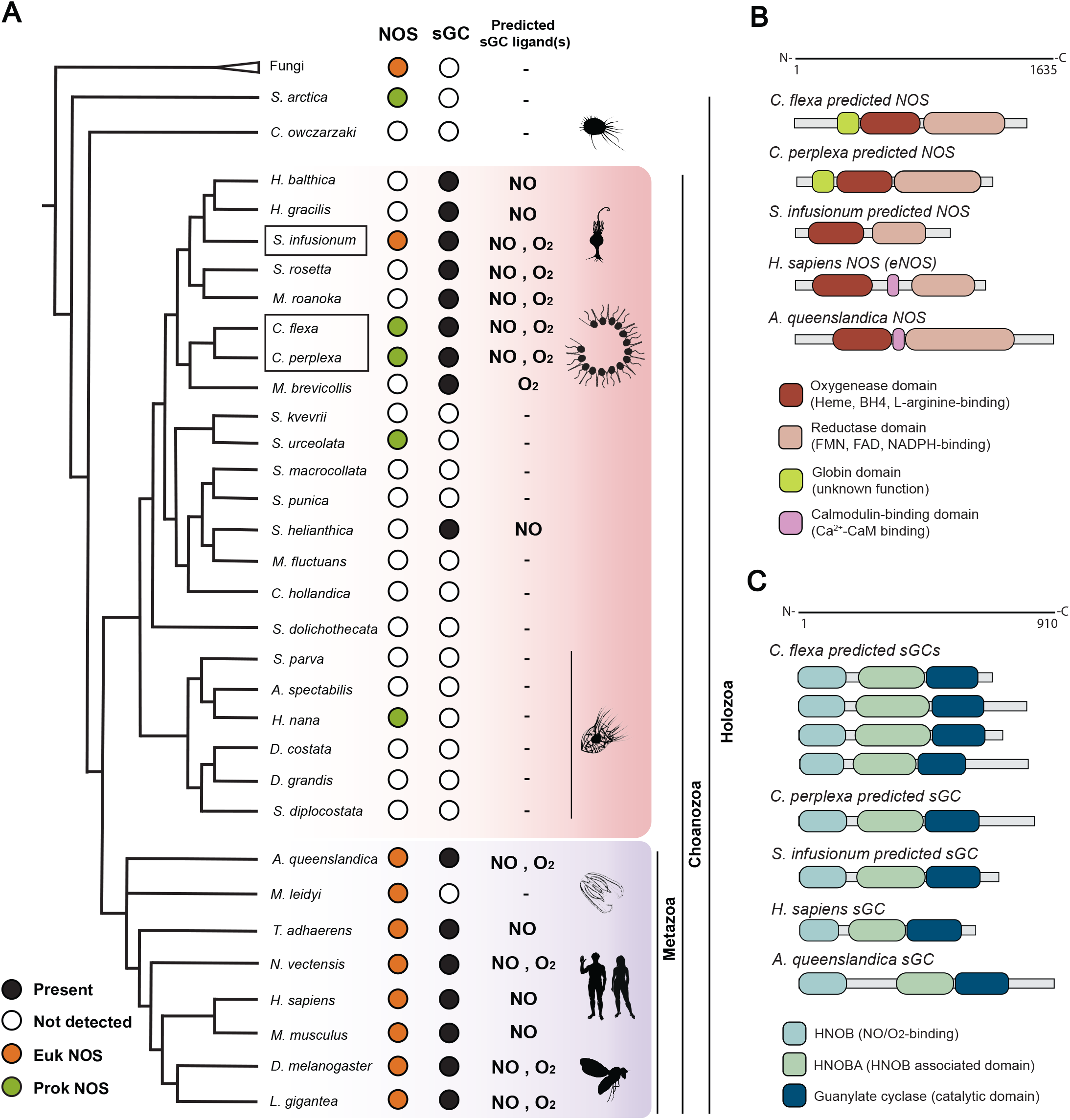
NO synthase (NOS) and soluble guanylate cyclases (sGC) predicted to bind either NO or O_2_ are broadly distributed across choanozoans, and all three are present in *C. flexa*. **(A)** Phylogenetic distribution of NOS and sGC across opisthokonts. *C. flexa*, its sister species *C. perplexa*, and *S. infusionum* encode both NOS and NO-sensitive sGCs, as do animals. **(B)** Choanoflagellate NOSs have the metazoan canonical oxygenase and reductase domain but lack the calcium-calmodulin binding domain. *C. flexa* and other choanoflagellate NOS encode an upstream globin domain with unknown function, also observed in cyanobacterial NOS. **(C)** *C. flexa* sGCs have the same domain architecture as animal sGCs.

Like metazoan NOS genes, choanoflagellate NOS genes encode the canonical oxygenase and reductase domains (Fig. 2B). Phylogenetic analysis revealed that one choanoflagellate NOS gene, that from *S. infusionum*, is closely related to those from metazoans and fungi (Fig. S1A). In contrast, all other choanoflagellate NOSs (including those from *C. flexa*) cluster with cyanobacterial NOSs, suggesting they might have been acquired through horizontal gene transfer from a cyanobacterial ancestor early in the evolution of choanoflagellates (Fig. S1A). These choanoflagellate NOS genes, like those of cyanobacteria, differ from metazoan NOS in that they encode an upstream globin domain with unknown function and lack the calmodulin-binding domain that mediates regulation of metazoan NOSs by Ca^2+^, suggesting that calcium signaling does not regulate NO synthesis in *C. flexa*. The *C. flexa* transcriptome also encodes complete biosynthetic pathways for the NOS cofactors BH_4_, FMN, FAD and NADPH (Fig. S2A) as well as downstream genes in the NO/cGMP signaling pathway: cGMP-dependent kinase (PKG), cGMP-gated ion channels (CNG) and cGMP-dependent phosphodiesterase (PDEG) (Fig. S2B), providing additional evidence that *C. flexa* employs NO signaling as part of its physiology.

Nearly all sGC genes from choanoflagellates, including *C. flexa*, encode the canonical domains observed in animal sGCs: the heme NO/O_2_-binding domain (HNOB or H-NOX), the HNOB associated domain (HNOBA) and the C-terminal catalytic domain (guanylate cyclase)^22^ (Fig. 2C). Phylogenetic analysis revealed that most choanoflagellate sGCs (including those of *C. flexa*) evolved from a single ancestral sGC found in the last common ancestor of choanoflagellates and metazoans which diversified separately into multiple paralogs in these two lineages (Fig. S1B). In contrast, the predicted sGC from one choanoflagellate species, *S. helianthica*, more closely resembles the sGCs of chlorophyte algae (which are the only protist group previously known to encode sGC proteins with a metazoan-like domain architecture^23^; Fig. S1B).

Importantly, not all animal sGCs are regulated by nitric oxide: in *Drosophila melanogaster* and *Caenorhabditis elegans*, so-called “atypical sGCs” preferentially bind soluble O_2_ instead of NO and are thought to be involved in the regulation of feeding by oxygen concentration^24-26^. Discrimination between NO and O_2_ is mediated by the presence of a hydrogen-bonding network, where a distal pocket tyrosine residue is critical for stabilizing the heme-O_2_ complex^27^. We generated preferential binding predictions based on this motif^27,28^ and found that both NO and O_2_-preferential binding sGCs are widely distributed among choanoflagellates with no obvious pattern (Fig. 2A; Fig. S2D). Interestingly, predicted NO-selective sGCs are present in choanoflagellate species in which NOS was not detected, suggesting that these species might detect NO from an exogenous source (i.e. environmental bacteria or other protists), might possess an alternative NO-producing mechanism, or might encode an NOS that was not detected in the transcriptome. In *C. flexa*, one out of four sGC transcripts was predicted to be selective for NO and was named Cf-sGC1 (Fig. S2D). The other two choanoflagellate species found to possess both a NOS and sGCs, *C. perplexa* and *S. infusionum*, were also predicted to encode at least one NO-sensitive sGC (Fig. 2A).

### NO/cGMP signaling controls colony contraction in *C. flexa*

To test whether NO signaling regulates collective contractions in *C. flexa*, we treated *C. flexa* cultures with several NO donors^29^, compounds capable of releasing NO in solution. We found that treatment of *C. flexa* with the NO donor proliNONOate led to an increase in intracellular NO as detected by the NO-sensitive fluorescent probe DAF-FM, demonstrating that it could be an effective reagent for studying NO signaling in *C. flexa in vivo* (Fig. 3A, S3A-C). Treatment of *C. flexa* with proliNONOate (Fig. 3B,C), DEANONOate (Fig. S3D) or NOC-12 (Fig. S3D) induced colony contraction within one to two minutes. Moreover, the inversion response of *C. flexa* to proliNONOate was concentration-dependent (Fig. 3D), reaching a plateau of nearly 100% inversion at a concentration of 0.1 μM. As a negative control, *C. flexa* did not invert in response to proline, the molecular backbone of proliNONOate and the end product of NO release (Fig. 3D). Taken together, these results indicate that NO is sufficient to induce colony contraction.

**Figure 3.**
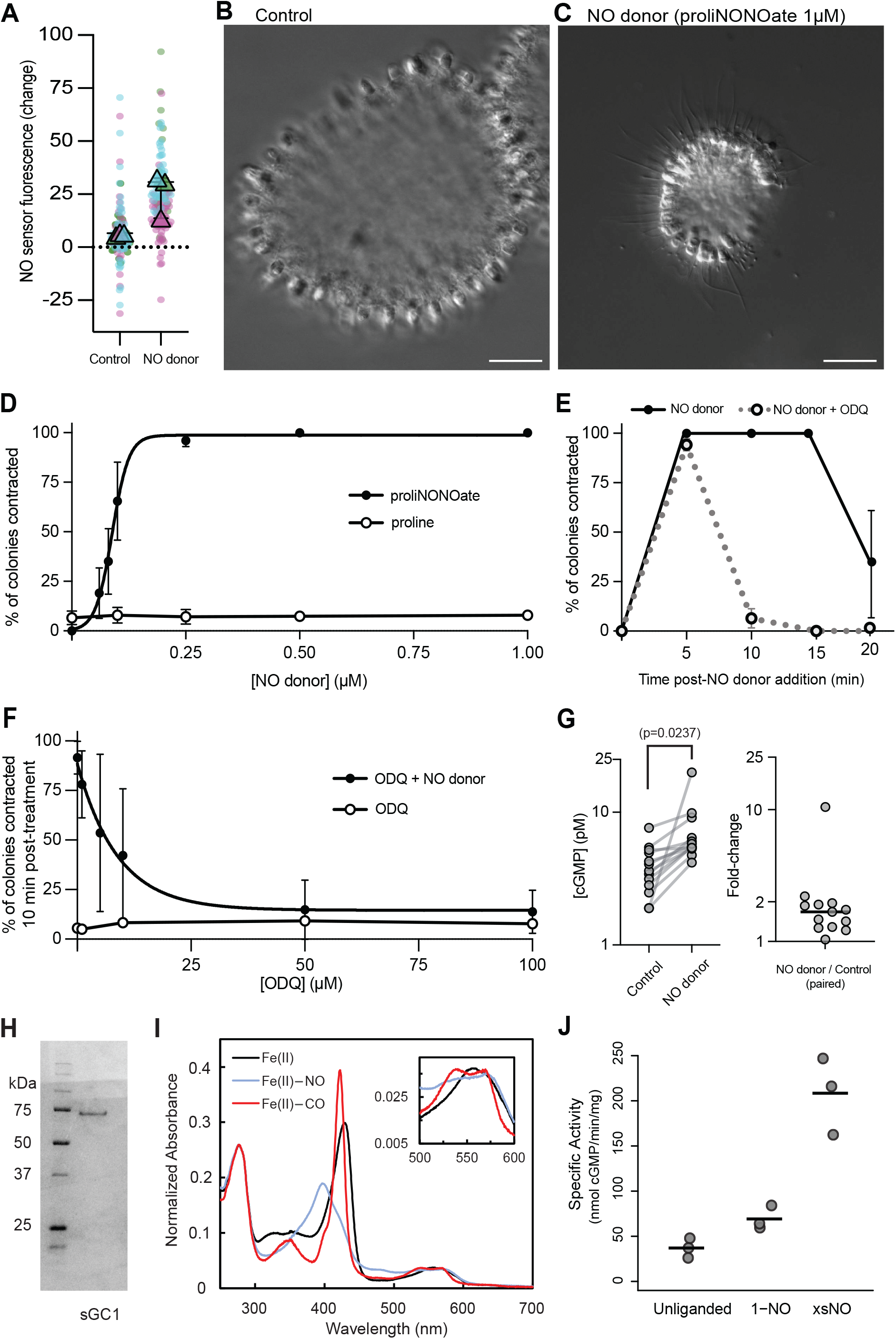
NO induced sustained contraction in *C. flexa* through sGC activation and cGMP synthesis. **(A)** NO release by proliNONOate was visualized by loading cells with the NO-sensitive fluorescent probe DAF-FM and measuring the change in intracellular fluorescence intensity over 30 minutes. Cells treated with NO donor showed a higher increase in fluorescence. Graph is a SuperPlot^67^ of N=3 biological replicates per condition, with sample sizes n=27, 33, 47 for the three control groups and n=26, 57, 42 for the three treated groups. *p=*0.0331 by an unpaired t-test. **(B-C)** NO induces colony contraction. While colonies treated with a negative control compound (proline) remained relaxed, colonies treated with 0.25 μM of the NO donor proliNONOate contracted within 1-2 minutes. **(D)** The NO donor proliNONOate induces contractions within ∼1 minute in a dose-dependent manner. A closely related molecule incapable of releasing NO (proline) had no effect over the same concentration range. **(E)** Inhibition of sGCs with 50 μM ODQ did not abolish NO-induced contraction at early time points but greatly reduced its duration. ODQ-treated colonies were contracted 5 minutes post-treatment with NO donor but relaxed soon after, while untreated colonies remain contracted for at least 5 more minutes. **(F)** Inhibition of sustained contraction by ODQ is dose-dependent. Colonies were incubated with different concentrations of ODQ for 1 hour before treatment with 0.25 μM of the NO donor proliNONOate. We quantified the percentage of contracted colonies 10 minutes after NO-treatment. In **(D-F)** each point represents the mean value of N=3 biological replicates with at least 30 colonies scored per biological replicate. Error bars are standard deviations. **(G)** Treatment of cells with a NO donor increased intracellular cGMP concentration almost 2-fold as quantified by ELISA. N=13 pairs of control/treated samples, *p*=0.024 by a paired T-test. **(H-J)** Purification, ligand binding properties and specific activity of Cf*-*sGC1. **(H)** Coomassie-stained SDS-PAGE gel of recombinant Cf*-*sGC1 expressed in *E. coli* and purified. Band in lane “sGC1” represents Cf*-*sGC1 with a monomeric molecular weight of 75.7 kDa. Left lane, molecular weight ladder. **(I)** UV-visible absorption spectra of *Cf* sGC1 under unliganded (black), NO-bound (blue), and CO-bound (red) conditions. Soret maxima: NO-bound: 429 nm; CO-bound: 423 nm; NO-bound: 399 nm. Inset, α/β bands show increased splitting upon ligand binding. **(J)** Specific activity of Cf-sGC1 under unliganded, equimolar quantities of NO (1-NO) and excess NO (xsNO) conditions. Initial rates were measured from activity assays performed at 25°C, pH 7.5 with 1.5 mM Mg^2+^-GTP substrate and 40 nM enzyme. Horizontal bars represent mean of three biological replicates.

We next investigated whether NO triggers cGMP synthesis in *C. flexa*. We found that a pan-inhibitor of sGCs, ODQ^30^, had no detectable effect on the percentage of contracted colonies 5 minutes after treatment with NO relative to cultures not treated with ODQ. However, a difference between the two conditions became evident 10 minutes after NO treatment, with nearly all ODQ-treated colonies relaxing into the uncontracted form, while all ODQ-untreated colonies remained contracted at least twice as long (Fig. 3E). The observation of rapid colony relaxation in ODQ-treated cultures was concentration-dependent (Fig. 3F).

These results demonstrate that the maintenance of NO-induced colony contraction requires sGC activity. If NO does indeed signal through the sGC/cGMP pathway, we predicted that treatment of colonies with an NO donor should increase intracellular cGMP concentration. We found that NO-treated cells contained consistently higher intracellular cGMP levels than untreated cells (∼2-fold increase on average, p=0.0237; Fig. 3G). Importantly, prior work has demonstrated that treatment of *C. flexa* with a cell-permeant form of cGMP (8-Br-cGMP) is sufficient to induce inversion^15^. These results further support the hypothesis that NO activates sGCs and stimulates the production of cGMP, which maintains *C. flexa* colony contraction.

### *C. flexa* sGC1 is an NO-selective and catalytically active component of the *C. flexa* NO/cGMP signaling pathway

Of the four sGCs expressed by *C. flexa*, only one (Cf-sGC1) was predicted to selective for NO (Fig. 1A). To directly test the existence of a canonical NO/sGC/cGMP pathway in *C. flexa*, we characterized ligand binding and activity of Cf-sGC1 that had been heterologously expressed in and purified from *E. coli* (Fig. 3H). The ligand specificity of sGCs can be determined by comparing the “Soret peak” maxima of the ultraviolet/visible (UV/vis) absorption after incubation with different gases, including NO, O_2_ or CO. The Fe(II), unliganded form of Cf-sGC1 exhibited a Soret peak at 429 nm and a single broad plateau in the α/β region (500-600 nm), consistent with that of an NO-selective sGC^28^ (Fig. 3I; table S1). In the presence of NO, the Soret peak shifted to 399 nm with two peaks in the α/β region, as characteristic for a 5-coordinate, high-spin NO-heme complex^28^ (Fig. 3I; Table S1). On the other hand, when exposed to atmospheric oxygen in the absence of NO, the UV-vis absorption spectrum of Cf-sGC1 did not change (Fig. 3I; Fig. S4), consistent with our prediction that Cf-sGC1 is an NO-selective sGC and does not bind O_2_. Moreover, like previously characterized animal sGCs, Cf-sGC1 also binds CO to form a 6-coordinate, low spin CO-heme complex^31^, evidenced by a Soret band maximum of 424 nm with two peaks in the α/β region (Fig. 3I; Table S1). Thus, the UV-vis spectroscopy results indicate that Cf-sGC1 binds diatomic gas ligands in a manner that resembles other well-characterized NO-selective sGCs.

In animals, NO-selective sGCs display three levels of activity: 1) in the absence of NO, the protein has a low basal guanylate cyclase activity; 2) when one NO molecule is bound at the heme moiety, the protein is partially activated (to several-fold the basal activity); and 3) in the presence of excess NO, the protein reaches maximal activation^32^. To characterize the enzymatic activity of Cf-sGC1, we measured cGMP production by purified Cf-sGC1 under unliganded, equimolar NO, and excess NO conditions using an endpoint activity assay. We found that Cf-sGC1 has an activity profile similar to that of animal sGCs, namely a 2-fold increase in activity in the equimolar NO condition, and ∼6-fold increase in the excess NO condition (Fig. 3J). Overall, Cf-sGC1 exhibits similar ligand binding properties and NO-stimulated activity profile to animal NO-specific sGCs^31^. These results further support the existence of NO/cGMP signaling in *C. flexa* and is consistent with it being mediated (at least in part) by Cf-sGC1.

### NO/cGMP signaling acts independently from most other inducers of colony contraction

In animals, NO signaling can be induced by a broad range of stimuli, which include chemical signals (for example, acetylcholine in mammalian blood vessels^33^), mechanical cues (for example, shear stress in blood vessels^34^), or heat shocks^35-38^. Interestingly, collar contractions in choanoflagellates can often be induced by mechanical stimuli, such as flow and touch^39-42^. We thus set out to test whether NO signaling in *C. flexa* responds to or intersects with environmental inducers of inversion.

We observed that *C. flexa* colonies invert in a matter of seconds in response to agitation of culture flasks (which presumably combines the effect of flow and shocks with other colonies or the walls of the flask) and to heat shocks (Fig. 4A,B). To test whether mechanically or heat-induced contraction requires NO/cGMP signaling, we incubated the colony cultures with the pan-sGC inhibitor ODQ and exposed them to either of the two different stressors. We found that inhibition of sGCs did not abolish mechanically induced or heat shock-induced contraction (Fig. 4A,B). Taken together, these findings suggest that the mechanosensitive, and thermosensitive pathways that induce inversion in *C. flexa* are independent of sGCs, and hint at complex behavioral regulation in this choanoflagellate.

**Figure 4.**
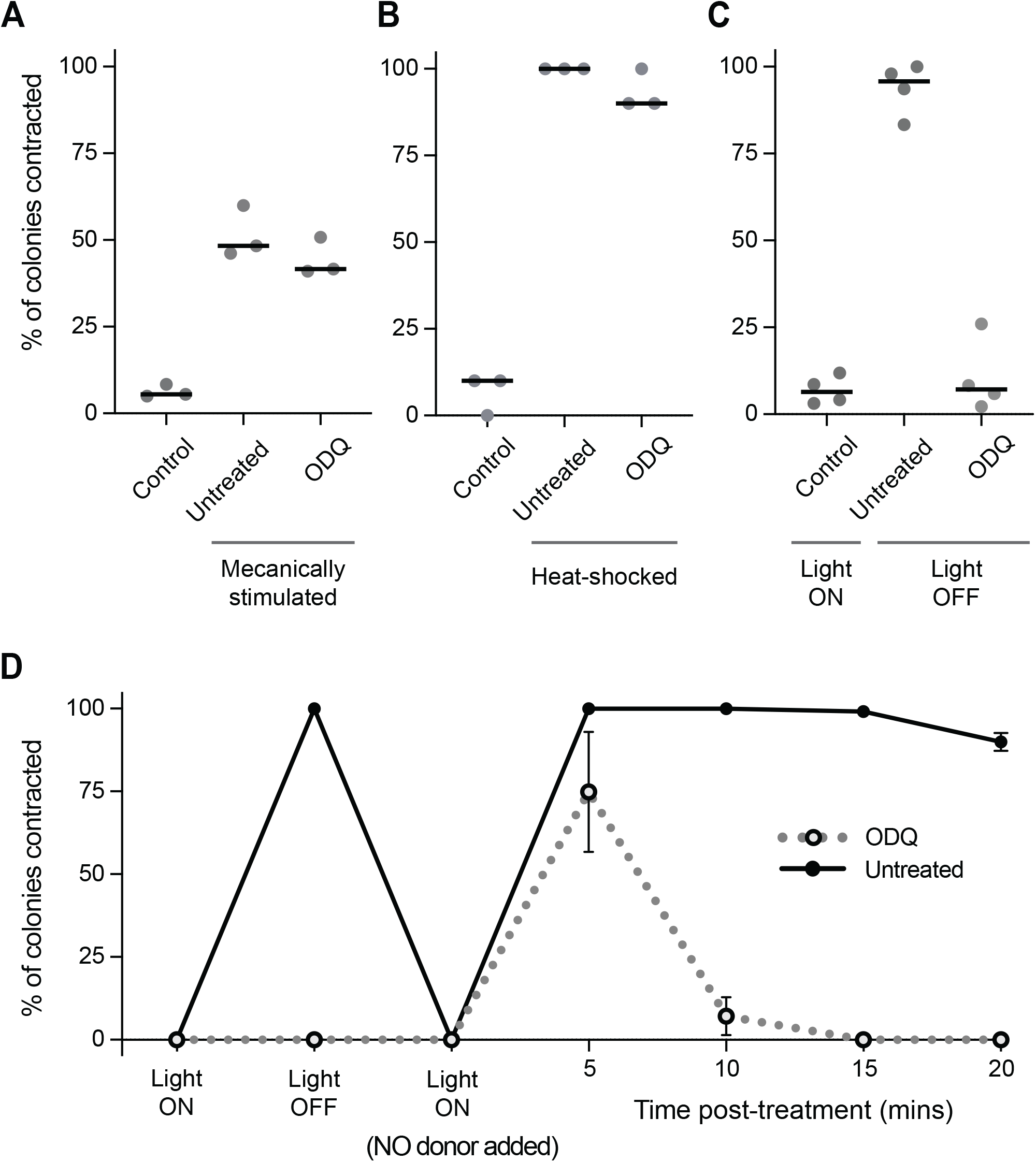
NO/cGMP acts independently of light-regulated, mechanically induced and heat shock-induced contractions in *C. flexa*. **(A-B)** Mechanical stimuli and heat-shock induce *C. flexa* colony contraction. Inhibition of sGC (50 μM ODQ) did not have an effect on mechanically or heat shock-induced contractions. **(C)** Inhibition of sGC (50 μM ODQ) abolished darkness-induced inversion (p<0.0001 by an unpaired t-test). **(D)** Colonies treated with ODQ did not respond to on/off changes in light but briefly contracted when exposed to NO donor, consistent with earlier results (Figure 3D-F). 10 minutes after treatment, ODQ-treated colonies were relaxed, while untreated colonies remained contracted for a longer period. In **(A-D)** each point is the average of N=3 **(A-B)** or N=4 **(C-D)** biological replicates with at least n=30 colonies scored per biological replicate. Error bars are standard deviations.

Previous work has shown that *C. flexa* colonies invert in response to light-to-dark transitions which they detect through a rhodopsin-cGMP pathway^15^. In the presence of light, a rhodopsin-phosphodiesterase hydrolyzes cGMP into GMP, thus preventing cGMP signaling. In darkness, the rhodopsin-phosphodiesterase is inactivated, which allows cGMP to accumulate and trigger colony inversion (Fig. 1B). Interestingly, this pathway requires the presence of cGMP, which is presumably synthesized by either particulate (i.e. membrane-bound) or soluble (i.e. cytosolic) guanylate cyclases^43^.

We next set out to answer whether sGCs are necessary for synthesizing the cGMP required for phototransduction. To address this, we treated light-sensitive colonies with ODQ and found that this entirely abolished darkness-induced inversion (Fig. 4C). These results suggest that sGCs (either NO-dependent or O_2_-dependent) are responsible for synthesizing baseline levels of cGMP that are then used during phototransduction. Even though ODQ-treated light-sensitive colonies did not invert in response to darkness, we confirmed that they could still respond to NO by undergoing brief contractions (which were sustained for a much shorter time than in controls; Fig. 4D), consistently with earlier results (Fig. 3E).

## DISCUSSION

Here we report the presence of NOS, sGCs and downstream components of NO/cGMP signaling in three choanoflagellate species, at least two of which (*C. flexa* and *C. perplexa*) are capable of collective contractions^15,44^. To our knowledge, this is the first observation of both NOS and sGCs in a non-animal. We found that NO causes sustained colony contraction in *C. flexa* and an increase in cGMP concentration in live cells, while inhibition of sGCs (and thereby cGMP concentrations) accelerated colony relaxation. Moreover, *in vitro* experiments confirmed that NO directly binds Cf-sGC1, which it activates with a two-step profile in response to NO concentration, as in animal sGCs^32^.

The observation that colonies treated with the sGC inhibitor initially contracted in response to NO at levels matching untreated colonies, only to relax much more quickly was unexpected. We hypothesize that NO-induced contractions are mediated through at least two different pathways: a slow pathway (described above) that maintains contraction and requires sGC/cGMP and an (unidentified) fast pathway independent of sGC/cGMP. Moreover, treatment of light-sensitive colonies with the sGC inhibitor abolished darkness-induced contractions (which are known to be mediated by cGMP^15^) but did not prevent NO-induced contractions, further supporting the existence of a second pathway. In other organisms, cGMP-independent NO signaling can involve S-nitrosation, the modification of proteins through the formation of an S-NO covalent bond^45,46^, although the direct targets and functions of S-nitrosation in animals are incompletely understood compared to NO/cGMP signaling. It is possible that the same mechanism explains the cGMP-independent pathway underlying NO-induced colony contraction in *C. flexa*.

The control of multicellular behavior by NO/cCMP signaling in *C. flexa* is reminiscent of its function in animals, most notably in sponges^47^. In the demosponges *Tethya wilhelma*^1^, *Ephydatia muelleri*^2^, and *Spongilla lacustris*^*3*^, NO induces global contractions and stops flagellar beating in choanocyte chambers, which interrupts feeding, allows expulsion of clumps of waste, and flushes the aquiferous canal system (a behavior sometimes called “sneezing”)^48^. Recently, single-cell RNA sequencing in *Spongilla lacustris* revealed that pinacocytes (epithelial cells that cover and shape the sponge body) co-express NOS and sGC^3,49^, the actomyosin contractility module and the transcription factor *Serum response facto*r (Srf), a master regulator of contractility^3,50,51^.

Control of motor and feeding behavior by NO/cGMP is also observed in cnidarians and some bilaterians. In the jellyfish *Aglantha digitale*, NO/cGMP signaling in neurons induces a switch from slow swimming (associated with feeding) to fast swimming (associated with escape) and inhibits tentacular ciliary beating^11^. In the sea pansy (a type of colonial cnidarian) *Renilla koellikeri*, NO/cGMP increases the amplitude of peristaltic contractions associated with the movement of body fluids through the gastrovascular cavity^17^. Finally, in the nudibranch *Clione limacine* and the snail *Lymnaea stagnalis*, NO activates both feeding and locomotory neural circuits^52-55^. Thus, as in *C. flexa*, the ancient functions of NO/cGMP signaling in animals may include the regulation of feeding and contraction^16,18,53,56-58^. Interestingly, NO signaling controls metamorphosis in sponges^7,59^, gastropods^60^, annelids^61^, echinoderms^62,63^ and ascidians^62,64,65^, thus regulating a switch from swimming to feeding during irreversible developmental programs.

In the future, identifying the function of Cf-NOS will require gene knock-out, which is not yet possible in *C. flexa*. Moreover, studies on *Trichoplax* (in which NO/cGMP signaling has been predicted to exist based on genomic data^66^), additional animal phyla, and other choanoflagellates will help flesh out reconstitutions of the early evolution of animal NO signaling.

## Acknowledgements

We thank Daniel J. Richter for sharing the *Ministeria* predicted proteome, and the whole King and Marletta labs for stimulating discussions. This work was supported by an EMBO long-term fellowship (ALTF 1474–2016) to TB, a Human Frontier Science Program long-term fellowship (000053/2017 L) to TB, the Institut Pasteur (G5 package; TB), the Howard Hughes Medical Institute (NK), the National Institute of Health (NIH R01GM127854; MM, YW, BG)), and the National Science Foundation Graduate Research Fellowship Program (Grant No. 1752814; JRR).

## Author contributions

JRR: Conceptualization, Methodology, Validation, Formal analysis, Investigation, Writing - Original draft, Writing - Review & editing, Visualization. YW: Conceptualization, Methodology, Validation, Formal analysis, Investigation, Writing - Original draft, Writing - Review & editing, Visualization. BGH: Conceptualization, Methodology. MM: Conceptualization, Methodology, Writing - Review & editing, Supervision, Project administration, Funding acquisition. NK: Conceptualization, Methodology, Writing - Review & editing, Supervision, Project administration, Funding acquisition. TB: Conceptualization, Methodology, Validation, Formal analysis, Investigation, Writing - Original draft, Writing - Review & editing, Supervision, Project administration, Funding acquisition.

## References

1. Ellwanger, K., and Nickel, M. (2006). Neuroactive substances specifically modulate rhythmic body contractions in the nerveless metazoon Tethya wilhelma (Demospongiae, Porifera). Front Zool 3, 7. 10.1186/1742-9994-3-7.

2. Elliott, G.R., and Leys, S.P. (2010). Evidence for glutamate, GABA and NO in coordinating behaviour in the sponge, Ephydatia muelleri (Demospongiae, Spongillidae). J Exp Biol 213, 2310–2321. 10.1242/jeb.039859.

3. Musser, J.M., Schippers, K.J., Nickel, M., Mizzon, G., Kohn, A.B., Pape, C., Ronchi, P., Papadopoulos, N., Tarashansky, A.J., Hammel, J.U., et al. (2021). Profiling cellular diversity in sponges informs animal cell type and nervous system evolution. Science 374, 717–723. 10.1126/science.abj2949.

4. Bogdan, C. (2015). Nitric oxide synthase in innate and adaptive immunity: an update. Trends Immunol 36, 161–178. 10.1016/j.it.2015.01.003.

5. Hillyer, J.F., and Estevez-Lao, T.Y. (2010). Nitric oxide is an essential component of the hemocyte-mediated mosquito immune response against bacteria. Dev Comp Immunol 34, 141–149. 10.1016/j.dci.2009.08.014.

6. Tomankova, S., Abaffy, P., and Sindelka, R. (2017). The role of nitric oxide during embryonic epidermis development of Xenopus laevis. Biol Open 6, 862–871. 10.1242/bio.023739.

7. Ueda, N., Richards, G.S., Degnan, B.M., Kranz, A., Adamska, M., Croll, R.P., and Degnan, S.M. (2016). An ancient role for nitric oxide in regulating the animal pelagobenthic life cycle: evidence from a marine sponge. Sci Rep 6, 37546. 10.1038/srep37546.

8. Gibbs, S.M., Becker, A., Hardy, R.W., and Truman, J.W. (2001). Soluble guanylate cyclase is required during development for visual system function in Drosophila. J Neurosci 21, 7705–7714.

9. Estephane, D., and Anctil, M. (2010). Retinoic acid and nitric oxide promote cell proliferation and differentially induce neuronal differentiation in vitro in the cnidarian Renilla koellikeri. Dev Neurobiol 70, 842–852. 10.1002/dneu.20824.

10. Pirtle, T.J., and Satterlie, R.A. (2021). Cyclic Guanosine Monophosphate Modulates Locomotor Acceleration Induced by Nitric Oxide but not Serotonin in Clione limacina Central Pattern Generator Swim Interneurons. Integr Org Biol 3, obaa045. 10.1093/iob/obaa045.

11. Moroz, L.L., Meech, R.W., Sweedler, J.V., and Mackie, G.O. (2004). Nitric oxide regulates swimming in the jellyfish Aglantha digitale. J Comp Neurol 471, 26–36. 10.1002/cne.20023.

12. Carr, M., Leadbeater, B.S., Hassan, R., Nelson, M., and Baldauf, S.L. (2008). Molecular phylogeny of choanoflagellates, the sister group to Metazoa. Proc Natl Acad Sci U S A 105, 16641–16646. 10.1073/pnas.0801667105.

13. Denninger, J.W., and Marletta, M.A. (1999). Guanylate cyclase and the .NO/cGMP signaling pathway. Biochim Biophys Acta 1411, 334–350. 10.1016/s0005-2728(99)00024-9.

14. Andreakis, N., D’Aniello, S., Albalat, R., Patti, F.P., Garcia-Fernandez, J., Procaccini, G., Sordino, P., and Palumbo, A. (2011). Evolution of the nitric oxide synthase family in metazoans. Mol Biol Evol 28, 163–179. 10.1093/molbev/msq179.

15. Brunet, T., Larson, B.T., Linden, T.A., Vermeij, M.J.A., McDonald, K., and King, N. (2019). Light-regulated collective contractility in a multicellular choanoflagellate. Science 366, 326–334. 10.1126/science.aay2346.

16. Colasanti, M., Venturini, G., Merante, A., Musci, G., and Lauro, G.M. (1997). Nitric oxide involvement in Hydra vulgaris very primitive olfactory-like system. J Neurosci 17, 493–499.

17. Anctil, M., Poulain, I., and Pelletier, C. (2005). Nitric oxide modulates peristaltic muscle activity associated with fluid circulation in the sea pansy Renilla koellikeri. J Exp Biol 208, 2005–2017. 10.1242/jeb.01607.

18. Colasanti, M., Persichini, T., and Venturini, G. (2010). Nitric oxide pathway in lower metazoans. Nitric Oxide 23, 94–100. 10.1016/j.niox.2010.05.286.

19. Robertson, H.M. (2009). The choanoflagellate Monosiga brevicollis karyotype revealed by the genome sequence: telomere-linked helicase genes resemble those of some fungi. Chromosome Res 17, 873–882. 10.1007/s10577-009-9078-2.

20. Fairclough, S.R., Chen, Z., Kramer, E., Zeng, Q., Young, S., Robertson, H.M., Begovic, E., Richter, D.J., Russ, C., Westbrook, M.J., et al. (2013). Premetazoan genome evolution and the regulation of cell differentiation in the choanoflagellate Salpingoeca rosetta. Genome Biol 14, R15. 10.1186/gb-2013-14-2-r15.

21. Richter, D.J., Fozouni, P., Eisen, M.B., and King, N. (2018). Gene family innovation, conservation and loss on the animal stem lineage. Elife 7. 10.7554/eLife.34226.

22. Derbyshire, E.R., and Marletta, M.A. (2009). Biochemistry of soluble guanylate cyclase. Handb Exp Pharmacol, 17–31. 10.1007/978-3-540-68964-5_2.

23. Horst, B.G., Stewart, E.M., Nazarian, A.A., and Marletta, M.A. (2019). Characterization of a Carbon Monoxide-Activated Soluble Guanylate Cyclase from Chlamydomonas reinhardtii. Biochemistry 58, 2250–2259. 10.1021/acs.biochem.9b00190.

24. Gray, J.M., Karow, D.S., Lu, H., Chang, A.J., Chang, J.S., Ellis, R.E., Marletta, M.A., and Bargmann, C.I. (2004). Oxygen sensation and social feeding mediated by a C. elegans guanylate cyclase homologue. Nature 430, 317–322. 10.1038/nature02714.

25. Huang, S.H., Rio, D.C., and Marletta, M.A. (2007). Ligand binding and inhibition of an oxygen-sensitive soluble guanylate cyclase, Gyc-88E, from Drosophila. Biochemistry 46, 15115–15122. 10.1021/bi701771r.

26. Cheung, B.H., Arellano-Carbajal, F., Rybicki, I., and de Bono, M. (2004). Soluble guanylate cyclases act in neurons exposed to the body fluid to promote C. elegans aggregation behavior. Curr Biol 14, 1105–1111. 10.1016/j.cub.2004.06.027.

27. Boon, E.M., and Marletta, M.A. (2005). Ligand discrimination in soluble guanylate cyclase and the H-NOX family of heme sensor proteins. Curr Opin Chem Biol 9, 441–446. 10.1016/j.cbpa.2005.08.015.

28. Boon, E.M., Huang, S.H., and Marletta, M.A. (2005). A molecular basis for NO selectivity in soluble guanylate cyclase. Nat Chem Biol 1, 53–59. 10.1038/nchembio704.

29. Cheng, J., He, K., Shen, Z., Zhang, G., Yu, Y., and Hu, J. (2019). Nitric Oxide (NO)-Releasing Macromolecules: Rational Design and Biomedical Applications. Front Chem 7, 530. 10.3389/fchem.2019.00530.

30. Zhao, Y., Brandish, P.E., Di Valentin, M., Schelvis, J.P., Babcock, G.T., and Marletta, M.A. (2000). Inhibition of soluble guanylate cyclase by ODQ. Biochemistry 39, 10848–10854. 10.1021/bi9929296.

31. Horst, B.G., Yokom, A.L., Rosenberg, D.J., Morris, K.L., Hammel, M., Hurley, J.H., and Marletta, M.A. (2019). Allosteric activation of the nitric oxide receptor soluble guanylate cyclase mapped by cryo-electron microscopy. Elife 8. 10.7554/eLife.50634.

32. Horst, B.G., and Marletta, M.A. (2018). Physiological activation and deactivation of soluble guanylate cyclase. Nitric Oxide 77, 65–74. 10.1016/j.niox.2018.04.011.

33. Doyle, M.P., and Duling, B.R. (1997). Acetylcholine induces conducted vasodilation by nitric oxide-dependent and -independent mechanisms. Am J Physiol 272, H1364–1371. 10.1152/ajpheart.1997.272.3.H1364.

34. Sriram, K., Laughlin, J.G., Rangamani, P., and Tartakovsky, D.M. (2016). Shear-Induced Nitric Oxide Production by Endothelial Cells. Biophys J 111, 208–221. 10.1016/j.bpj.2016.05.034.

35. Zhang, L., Liu, Q., Yuan, X., Wang, T., Luo, S., Lei, H., and Xia, Y. (2013). Requirement of heat shock protein 70 for inducible nitric oxide synthase induction. Cell Signal 25, 1310–1317. 10.1016/j.cellsig.2013.02.004.

36. Dulce, R.A., Mayo, V., Rangel, E.B., Balkan, W., and Hare, J.M. (2015). Interaction between neuronal nitric oxide synthase signaling and temperature influences sarcoplasmic reticulum calcium leak: role of nitroso-redox balance. Circ Res 116, 46–55. 10.1161/CIRCRESAHA.116.305172.

37. Rai, K.K., Pandey, N., and Rai, S.P. (2020). Salicylic acid and nitric oxide signaling in plant heat stress. Physiol Plant 168, 241–255. 10.1111/ppl.12958.

38. Giovine, M., Pozzolini, M., Favre, A., Bavestrello, G., Cerrano, C., Ottaviani, F., Chiarantini, L., Cerasi, A., Cangiotti, M., Zocchi, E., et al. (2001). Heat stress-activated, calcium-dependent nitric oxide synthase in sponges. Nitric Oxide 5, 427–431. 10.1006/niox.2001.0366.

39. Leadbeater, B.S. (1983). Distribution and chemistry of microfilaments in choanoflagellates, with special reference to the collar and other tentacle systems. Protistologica 19, 157–166.

40. James-Clark, H. (1868). XXII.—On the Spongiæ ciliatæ as Infusoria flagellata; or observations on the structure, animality, and relationship of Leucosolenia botryoides, Bowerbank. Annals and Magazine of Natural History 1, 133–142. 10.1080/00222936808695657.

41. Nguyen, N.M., Merle, T., Broders, F., Brunet, A.-C., Sarron, F., Jha, A., Genisson, J.-L., Rottinger, E., and Farge, E. (2020). Evolutionary Emergence of First Animal Organisms Triggered by Environmental Mechano-Biochemical Marine Stimulation. bioRxiv, 2020.2012.2003.407668. 10.1101/2020.12.03.407668.

42. Andrews, G.F. (1897). The living substance as such and as organism. Supplement to Journal of Morphology XII N. 2.

43. Zhang, X., and Cote, R.H. (2005). cGMP signaling in vertebrate retinal photoreceptor cells. Front Biosci 10, 1191–1204. 10.2741/1612.

44. Leadbeater, B.S.C. (1983). Life-history and ultrastructure of a new marine species of Proterospongia (Choanoflagellida). Journal of the Marine Biological Association of the United Kingdom 63, 135 – 160. 10.1017/S0025315400049857.

45. Broniowska, K.A., Diers, A.R., and Hogg, N. (2013). S-nitrosoglutathione. Biochim Biophys Acta 1830, 3173–3181. 10.1016/j.bbagen.2013.02.004.

46. Smith, B.C., and Marletta, M.A. (2012). Mechanisms of S-nitrosothiol formation and selectivity in nitric oxide signaling. Curr Opin Chem Biol 16, 498–506. 10.1016/j.cbpa.2012.10.016.

47. Simion, P., Philippe, H., Baurain, D., Jager, M., Richter, D.J., Di Franco, A., Roure, B., Satoh, N., Queinnec, E., Ereskovsky, A., et al. (2017). A Large and Consistent Phylogenomic Dataset Supports Sponges as the Sister Group to All Other Animals. Curr Biol 27, 958–967. 10.1016/j.cub.2017.02.031.

48. Elliott, G.R., and Leys, S.P. (2007). Coordinated contractions effectively expel water from the aquiferous system of a freshwater sponge. J Exp Biol 210, 3736–3748. 10.1242/jeb.003392.

49. Nickel, M., Scheer, C., Hammel, J.U., Herzen, J., and Beckmann, F. (2011). The contractile sponge epithelium sensu lato--body contraction of the demosponge Tethya wilhelma is mediated by the pinacoderm. J Exp Biol 214, 1692–1698. 10.1242/jeb.049148.

50. Miano, J.M., Long, X., and Fujiwara, K. (2007). Serum response factor: master regulator of the actin cytoskeleton and contractile apparatus. Am J Physiol Cell Physiol 292, C70–81. 10.1152/ajpcell.00386.2006.

51. Brunet, T., Fischer, A.H., Steinmetz, P.R., Lauri, A., Bertucci, P., and Arendt, D. (2016). The evolutionary origin of bilaterian smooth and striated myocytes. Elife 5. 10.7554/eLife.19607.

52. Moroz, L.L., Norekian, T.P., Pirtle, T.J., Robertson, K.J., and Satterlie, R.A. (2000). Distribution of NADPH-diaphorase reactivity and effects of nitric oxide on feeding and locomotory circuitry in the pteropod mollusc, Clione limacina. J Comp Neurol 427, 274–284.

53. Moroz, L.L., and Kohn, A.B. (2011). Parallel evolution of nitric oxide signaling: diversity of synthesis and memory pathways. Front Biosci (Landmark Ed) 16, 2008–2051. 10.2741/3837.

54. Moroz, L.L., Park, J.H., and Winlow, W. (1993). Nitric oxide activates buccal motor patterns in Lymnaea stagnalis. Neuroreport 4, 643–646. 10.1097/00001756-199306000-00010.

55. Kobayashi, S., Sadamoto, H., Ogawa, H., Kitamura, Y., Oka, K., Tanishita, K., and Ito, E. (2000). Nitric oxide generation around buccal ganglia accompanying feeding behavior in the pond snail, Lymnaea stagnalis. Neurosci Res 38, 27–34. 10.1016/s0168-0102(00)00136-x.

56. Yabumoto, T., Takanashi, F., Kirino, Y., and Watanabe, S. (2008). Nitric oxide is involved in appetitive but not aversive olfactory learning in the land mollusk Limax valentianus. Learn Mem 15, 229–232. 10.1101/lm.936508.

57. Cristino, L., Guglielmotti, V., Cotugno, A., Musio, C., and Santillo, S. (2008). Nitric oxide signaling pathways at neural level in invertebrates: functional implications in cnidarians. Brain Res 1225, 17–25. 10.1016/j.brainres.2008.04.056.

58. Jacklet, J.W. (1997). Nitric oxide signaling in invertebrates. Invert Neurosci 3, 1–14. 10.1007/BF02481710.

59. Song, H., Hewitt, O.H., and Degnan, S.M. (2021). Arginine Biosynthesis by a Bacterial Symbiont Enables Nitric Oxide Production and Facilitates Larval Settlement in the Marine-Sponge Host. Curr Biol 31, 433–437 e433. 10.1016/j.cub.2020.10.051.

60. Froggett, S.J., and Leise, E.M. (1999). Metamorphosis in the Marine Snail Ilyanassa obsoleta, Yes or NO? Biol Bull 196, 57–62. 10.2307/1543167.

61. Biggers, W.J., Pires, A., Pechenik, J.A., Johns, E., Patel, P., Polson, T., and Polson, J. (2012). Inhibitors of nitric oxide synthase induce larval settlement and metamorphosis of the polychaete annelidCapitella teleta. Invertebrate Reproduction & Development 56, 1–13. 10.1080/07924259.2011.588006.

62. Bishop, C.D., and Brandhorst, B.P. (2003). On nitric oxide signaling, metamorphosis, and the evolution of biphasic life cycles. Evol Dev 5, 542–550. 10.1046/j.1525-142x.2003.03059.x.

63. Bishop, C.D., and Brandhorst, B.P. (2007). Development of nitric oxide synthase-defined neurons in the sea urchin larval ciliary band and evidence for a chemosensory function during metamorphosis. Dev Dyn 236, 1535–1546. 10.1002/dvdy.21161.

64. Ueda, N., and Degnan, S.M. (2013). Nitric oxide acts as a positive regulator to induce metamorphosis of the ascidian Herdmania momus. PLoS One 8, e72797. 10.1371/journal.pone.0072797.

65. Comes, S., Locascio, A., Silvestre, F., d’Ischia, M., Russo, G.L., Tosti, E., Branno, M., and Palumbo, A. (2007). Regulatory roles of nitric oxide during larval development and metamorphosis in Ciona intestinalis. Dev Biol 306, 772–784. 10.1016/j.ydbio.2007.04.016.

66. Moroz, L.L., Romanova, D.Y., Nikitin, M.A., Sohn, D., Kohn, A.B., Neveu, E., Varoqueaux, F., and Fasshauer, D. (2020). The diversification and lineage-specific expansion of nitric oxide signaling in Placozoa: insights in the evolution of gaseous transmission. Sci Rep 10, 13020. 10.1038/s41598-020-69851-w.

67. Lord, S.J., Velle, K.B., Mullins, R.D., and Fritz-Laylin, L.K. (2020). SuperPlots: Communicating reproducibility and variability in cell biology. J Cell Biol 219. 10.1083/jcb.202001064.

68. Abbas, C.A., and Sibirny, A.A. (2011). Genetic control of biosynthesis and transport of riboflavin and flavin nucleotides and construction of robust biotechnological producers. Microbiol Mol Biol Rev 75, 321–360. 10.1128/MMBR.00030-10.

69. Zhao, Y., Brandish, P.E., Ballou, D.P., and Marletta, M.A. (1999). A molecular basis for nitric oxide sensing by soluble guanylate cyclase. Proc Natl Acad Sci U S A 96, 14753–14758. 10.1073/pnas.96.26.14753.

70. Torruella, G., de Mendoza, A., Grau-Bove, X., Anto, M., Chaplin, M.A., del Campo, J., Eme, L., Perez-Cordon, G., Whipps, C.M., Nichols, K.M., et al. (2015). Phylogenomics Reveals Convergent Evolution of Lifestyles in Close Relatives of Animals and Fungi. Curr Biol 25, 2404–2410. 10.1016/j.cub.2015.07.053.

71. Dereeper, A., Guignon, V., Blanc, G., Audic, S., Buffet, S., Chevenet, F., Dufayard, J.F., Guindon, S., Lefort, V., Lescot, M., et al. (2008). Phylogeny.fr: robust phylogenetic analysis for the non-specialist. Nucleic Acids Res 36, W465–469. 10.1093/nar/gkn180.

72. Letunic, I., and Bork, P. (2021). Interactive Tree Of Life (iTOL) v5: an online tool for phylogenetic tree display and annotation. Nucleic Acids Res 49, W293–W296. 10.1093/nar/gkab301.

73. Barr, I., and Guo, F. (2015). Pyridine Hemochromagen Assay for Determining the Concentration of Heme in Purified Protein Solutions. Bio Protoc 5. 10.21769/bioprotoc.1594.

