## Supplementary material for "Nitric oxide signaling controls collective contractions in a colonial choanoflagellate"

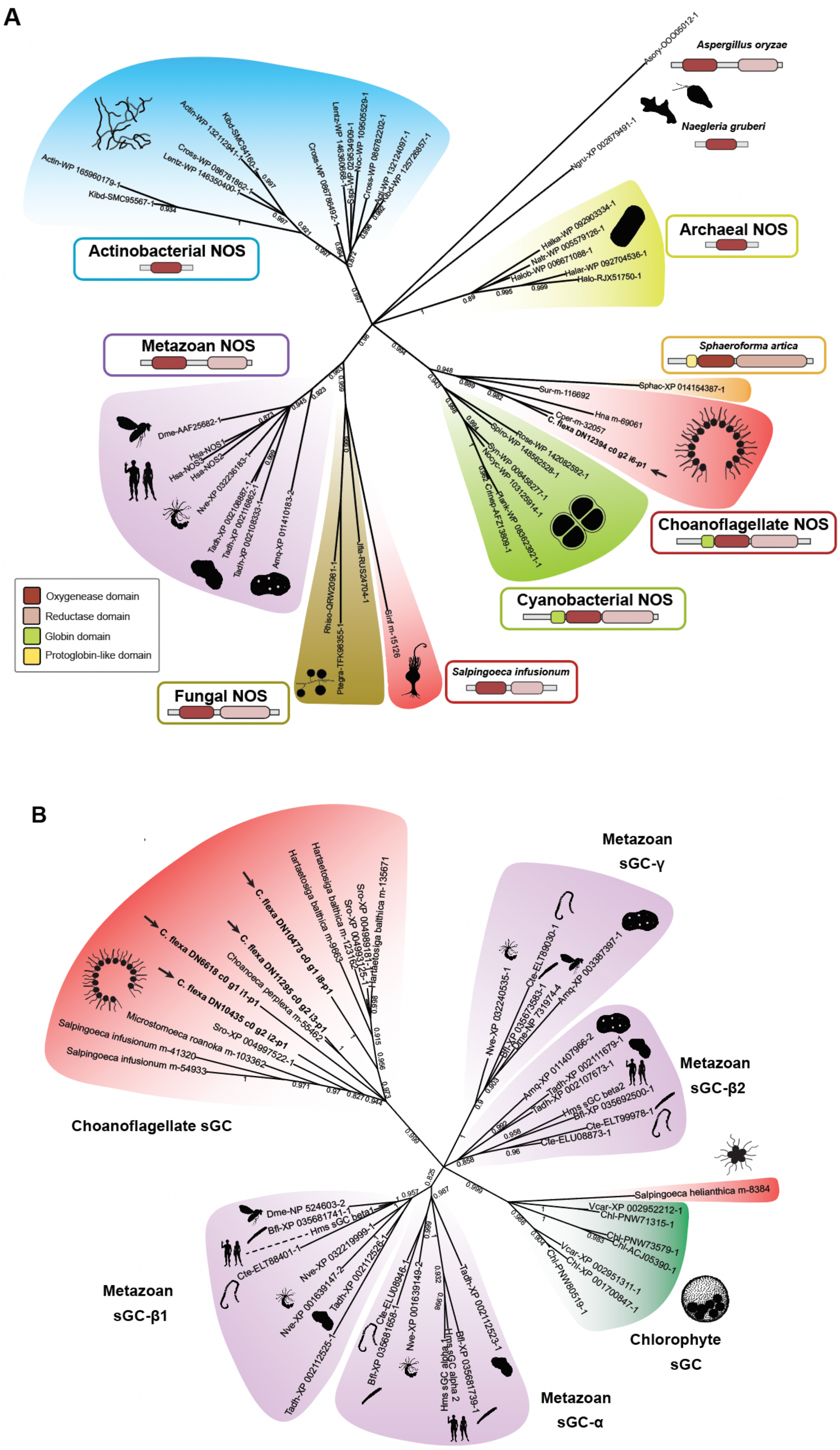
A
Actinobacterial NOS
Aspergillus oryzae
Naegleria gruberi
Archaeal NOS
Metazoan NOS
Sphaeroforma artica
Choanoflagellate NOS
Cyanobacterial NOS
Oxygenease domain
Reductase domain
Globin domain
Protoglobin-like domain
Fungal NOS
Salpingoeca infusionum
B
Metazoan
sGC-γ
Metazoan
sGC-β2
Choanoflagellate sGC
Chlorophyte
sGC
Metazoan
sGC-β1
Metazoan
sGC-α

**Figure S1.** Phylogenetic tree of NOS and sGC. **(A)** Phylogenetic tree of NOS from a selection of fully sequenced eukaryotic, archaeal, and bacterial genomes. Alignments and phylogenetic reconstruction were performed on the oxygenase domain only. The clades recovered tend to share common domain architectures out of the oxygenase domain (such as the reductase domain in eukaryotic NOSs and the globin domain in choanoflagellate and cyanobacterial NOSs), providing independent support to the phylogeny. The sister-group relationship between choanoflagellate and cyanobacterial NOSs as well as the shared domain architecture suggest a history of horizontal gene transfer. See Material and Methods for species name abbreviations. **(B)** Phylogenetic tree of sGC from a selection of fully sequenced animal and chlorophyte genomes, and choanoflagellate genomes and transcriptomes. Analysis was performed based on the full protein sequence. See Material and Methods for species name abbreviations.

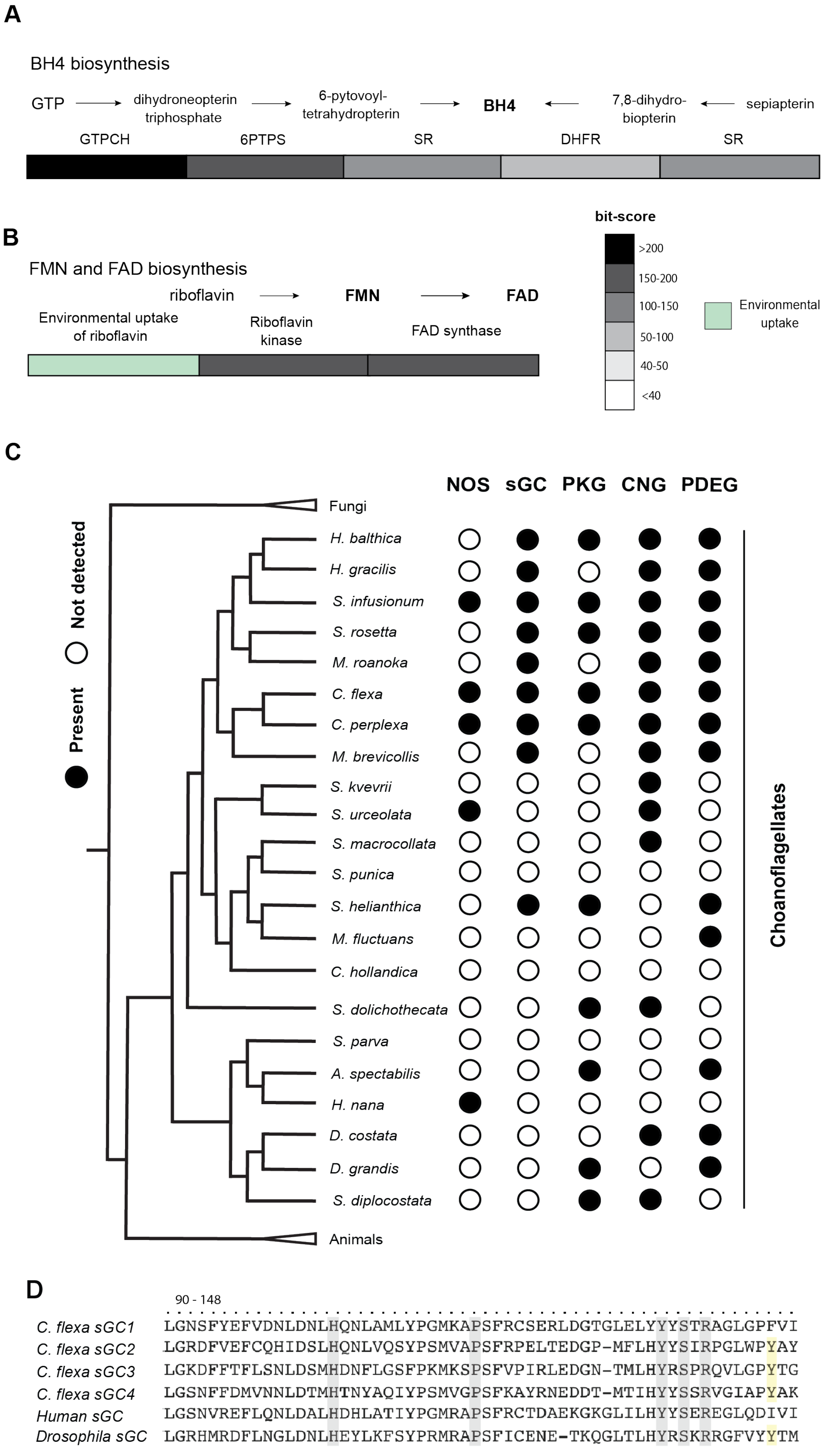
A
BH4 biosynthesis
GTP
dihydroneopterin triphosphate
6-pytovoyl-tetrahydropterin
BH4
7,8-dihydro-biopterin
sepiapterin
GTPCH
6PTPS
SR
DHFR
SR
B
FMN and FAD biosynthesis
riboflavin
FMN
FAD
Environmental uptake of riboflavin
Riboflavin kinase
FAD synthase
bit-score
>200
150-200
100-150
50-100
40-50
<40
Environmental uptake
C
NOS
sGC
PKG
CNG
PDEG
Present
Not detected
Fungi
H. balthica
H. gracilis
S. infusionum
S. rosetta
M. roanoka
C. flexa
C. perplexa
M. brevicollis
S. kvevrii
S. urceolata
S. macrocollata
S. punica
S. helianthica
M. fluctuans
C. hollandica
S. dolichothecata
S. parva
A. spectabilis
H. nana
D. costata
D. grandis
S. diplocostata
Animals
Choanoflagellates
D
90 - 148
C. flexa sGC1 LGNSFYEFVDNLDNLHQNLAMLYPGMKAPSFRCSERLDGTGLELYYYSTRAGLGPFVI
C. flexa sGC2 LGRDFVEFCQHIDSLHQNLVQSYPSMVAPSFRPELTEDGP-MFLHYYSIRPGLWPYAY
C. flexa sGC3 LGKDFFTFLSNLDSMHDNFLGSFPKMKSPSFVPIRLEDGN-TMLHYRSPRQVLGPYTG
C. flexa sGC4 LGSNFFDMVNNLDTMHHTNYAQIYPSMVGPSFKAYRNEDDT-MTIHYYSSRVGIAPYAK
Human sGC LGSNVREFLQNLDALHDHLATIYPGMRAPSFRCTDAEKGKGLILHYYSEREGLQDIVI
Drosophila sGC LGRHMRDFLNGLDNLHEYLKFSYPRMRAPSFICENE-TKQGLTLHYRSKRRGFVYYTM

**Figure S2.** *C. flexa* encodes the complete biosynthetic pathways for the NOS cofactors and downstream NO/cGMP signaling components. **(A-B)** The *C. flexa* transcriptome were searched for genes encoding enzymes in the tetrahydrobiopterin (BH4; **A**) and FMN/FAD **(B)** biosynthesis pathways using BLASTP. For each step in the pathway, multiple bacterial, plant, fungal and/or animal genes were used as queries (see Methods), and the highest returned bit-score is shown. *C. flexa* encodes the complete synthetic pathway. Riboflavin is assumed to be uptaken from the environment[68]. **(C)** Phylogenetic distribution of NO/cGMP signaling downstream components. In animals, cGMP is known to signal through cGMP-dependent kinases (PKG), cGMP-gated ion channels (CNG) and cGMP-dependent phosphodiesterases (PDEG). The three choanoflagellate species detected to express a NOS and sGC (*C. flexa*, *C. perplexa* and *S. infusionum*) also express PKG, CNG and PDEG. **(D)** We made preferential binding predictions for *C. flexa* sGCs based on NO-sensitive and $O_2$-binding metazoan sGCs[25,27,28,69]. *C. flexa* sGC partial alignment with human sGC (NO-sensitive) and *Drosophila* sGC ($O_2$-binding). Amino acids important for heme binding are highlighted in grey. A distal tyrosine residue is highlighted in yellow, previously shown to be highly important for $O_2$ binding[27]. We predicted one NO-sensitive (sGC1) and three $O_2$-binding sGCs (sGC2, sGC3, sGC4).

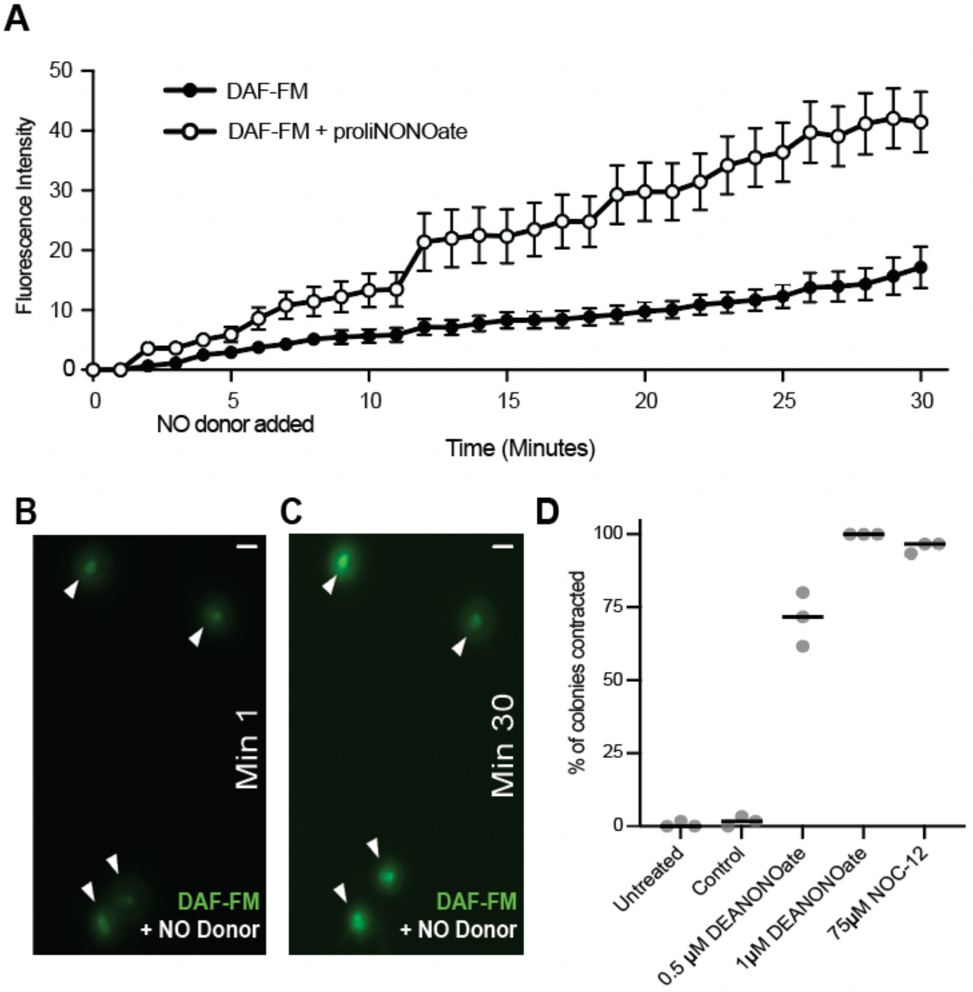


**Figure S3.** Treatment with NO donors increase intracellular NO levels and induce colony contraction. **(A-C)** Intracellular NO levels over time. We labeled NO with the fluorescent probe DAF-FM and measured fluoresce intensity every minute for 30 minutes. NO donor was added at minute five. Intracellular fluorescence in NO donor-treated cells display a higher increase over time compared to control. Small increase in fluorescence in control group may be explained by unwashed dye incorporating inside of the cells and/or basal physiological NO levels. Intracellular intensity was measured using Image J software. **(B-C)** Representative micrographs are shown on the right. White arrows are pointing individual cells. Scale bar: 5 µM. NO donor was added at minute five. **(D)** NO donors DEANONOate and NOC-12 also induce colony contraction.

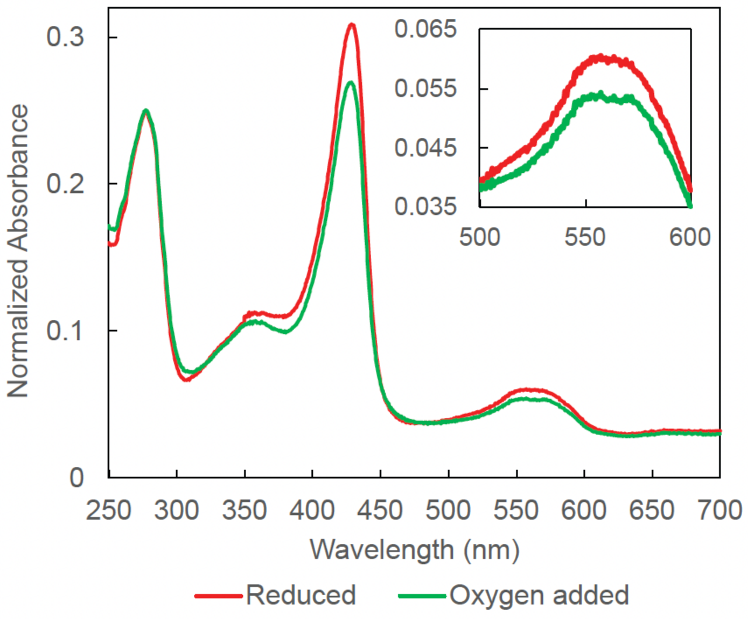


**Figure S4.** The UV-vis spectrum of Cf-sGC1 upon exposure to $O_2$, normalized to $A_{280}$. The Soret to $A_{280}$ ratio decreased when oxygen was added to the sample, which could indicate oxidation of the heme and loss of the heme through the process. There was no observable formation of a low-spin 6-coordinate Fe(II)-$O_2$ complex (as indicated by the lack of shift of the Soret peak).

**Table S1.** *Cf* sGC1 UV-vis absorption wavelengths. *Rattus norvegicus* sGC is a well-characterized, NO-selective sGC.

| Protein | Condition | Soret (nm) | α band (nm) | β band (nm) | References |
|---|---|---|---|---|---|
| ***Cf* sGC1** | Unliganded | 429 | 559 | | This work |
| | NO | 399 | 572 | 540 | |
| | CO | 423 | 568 | 538 | |
| ***Rattus norvegicus* sGC** | Unliganded | 431 | 562 | | (Zhao 1997) |
| | NO | 399 | 572 | 537 | |
| | CO | 423 | 568 | 541 | |

### Materials and methods

#### Culture of *Choanoeca flexa*

Colonies were cultured in 1% to 15% Cereal Grass Medium (CGM3) in artificial seawater (ASW). Polyxenic cultures (continuously passaged from a previously described environmental isolate[15]) were maintained at 22°C under a light-dark cycle of 12:12 hours in a Caron low temperature incubator equipped with a lamp (Venoya Full Spectrum 150W Plant Growth LED) controlled by a programmable timer (Leviton VPT24-1PZ Vizia). Polyxenic cultures used in most experiments were not light-sensitive, possibly due to progressive loss of bacterial diversity during serial passaging (as bacterially provided retinal is known to be required for photosensation in *C. flexa*[15]). Light-sensitive sheets used in photosensation experiments (Fig. 4C,D) were thawed from stocks that had been frozen immediately after clonal isolation from a Curaçao isolate and cultured as described above.

#### Light microscopy – Imaging

Colonies were imaged in FluoroDishes (World Precision Instruments FD35-100) by differential interference contrast (DIC) microscopy using a 20x Zeiss objective mounted on a Zeiss Observer Z.1 with Hamamatsu Orca Flash 4.0 V2 CMOS camera (C1140-22CU).

#### Small molecule compound treatments and colony inversion assays

Small molecule inhibitor treatments and colony inversion assays were performed in 24-well plates (Fischer Scientific 09-761-146) containing 1 mL *C. flexa* culture per well. ODQ (pan-soluble guanylate cyclase inhibitor) was added 1 hour before behavioral assays. Addition of each small molecule compound was followed by a gentle swirl of the 24-well plate to ensure mixing. For each assay, all colonies visible within a well were counted (at least 30 colonies per biological replicate).

*NO donor-induced inversion:* The NO donors proliNONOate (Cayman Chemical Company 82145) and DEA-NONOate (Cayman Chemical Company 82145) were dissolved according to provider's instructions and stored as single-use aliquots at -80°C. Addition of NO donors induced inversion within 1-2 minutes. Prior to counting, colonies were fixed by addition of 16% ice-cold PFA in a 1:3 volumetric ratio, resulting in a final concentration of 4% PFA. Contracted and relaxed colonies were then manually counted by observation under a Leica DMIL LED transmitted light microscope.

*Light-induced sheet inversion*: After treatment with small molecule compounds, light-to-dark transitions were performed by manually switching off the light source of the DMIL LED microscope. The "light off" condition lasted for one minute before sheets were fixed and scored as described above.

*Mechanically induced sheet inversion*: 3 mL of *C. flexa* culture were transferred to T12.5 culture flasks (Fisher Scientific 353107) and mechanically stimulated by vortexing on a Vortex Genie 2 (Scientific industries) on "Slow" setting for 5 seconds. Sheets were immediately fixed and scored as above.

*Heat shock-induced sheet inversion*: colonies in 24-well plates were treated with inhibitors as described above and placed at the surface of a 37°C warm bath for one minute. Sheets were immediately fixed and scored as above.

**cGMP ELISA**

For *in vivo* quantification of cGMP, we used an ENZO Direct cGMP ELISA kit (ADI-900-014, 96 wells) as directed by the manufacturer. For each biological replicate, 90 mL of dense (>$10^6$ cells/mL) *C. flexa* culture was centrifuged for 5 minutes at 3000 x g and resuspended in 25 mL of ASW to wash the bacteria away. After the third wash, the cells were resuspended in 200 µL of ASW and split into one control (100 µL) and one treated sample (100 µL). The samples were lysed and quantified in parallel in each assay. Colonies from the "NO donor" group were treated with 0.25 µM proliNONOate 5 minutes before lysis. Values were read on a SpectraMax M3 plate reader (Molecular Devices).

**NO labeling, imaging, and image analysis**

*C. flexa* cultures were transferred into 15 mL Falcon tubes and vortexed in "Fast" setting on a Vortex Genie 2 for one minute to dissociate colonies into single cells. Cells were washed 3 times with artificial seawater (ASW) by centrifuging them for 5 minutes at 3000 x g and resuspending them in 25 mL of ASW. After the last wash, cells were resuspended in 1.5 mL ASW and transferred into an 1.5 mL Eppendorf tube. Cells were incubated in 10µM DAF-FM (Invitrogen, D-23844) for 1 hour and rinsed twice with ASW to wash away the unincorporated dye. Cells were then transferred into a FluoroDish charged with a Corona surface treater and coated with poly-D-lysine, following a previously published protocol[15]. We let the cells adhere to the bottom of the dish for 30 minutes before imaging on a Z.1 Zeiss Imager with a Hamamatsu Orca Flash 4.0 V2 CMOS camera (C11440-22CU) and a 40x water immersion objective (C-Apochromat, 1.1 NA) for DIC and green epifluorescence imaging with a frame rate of 1 frame per minute. The NO donor (0.25µM proliNONOate) was added 5 minutes after imaging began. We quantified intracellular fluorescence intensity using ImageJ. Change in fluorescence intensity was calculated by substracting the fluorescence intensity at minute 1 from fluorescence intensity at minute 30.

**Phylogenetic analysis and protein domain identification**

We screened a selection of fully sequenced genomes for homologs of sGC and NOS with the following strategy: the protein sequences of *Homo sapiens* sGC-α1 and brain nitric oxide synthase (NOS1) were used as BLASTp queries against the NCBI database restricted to the following list of species:

- Eukaryotes: *Homo sapiens* (Hsa), *Branchiostoma floridae* (Bfl), *Drosophila melanogaster* (Dme), *Capitella teleta* (Cte), *Nematostella vectensis* (Nve), *Amphimedon queeslandica* (Amq), *Mnemiopsis leidyi, Trichoplax adhaerens* (Tadh), *Salpingoeca rosetta* (Sro), *Capsaspora owczarzaki, Sphaereoforma arctica* (Sphac), *Abeoforma whisleri, Creolimax fragrantissima, Pirum gemmata, Aspergillus oryzae* (Asory), *Jimgerdemannia flammicorona* (Jifla), *Rhizoctonia solani* (Rhiso), *Pterula gracilis* (Ptegra), *Schizosaccharomyces pombe, Tuber melanosporum, Cryptococcus neoformans, Ustilago maydis, Cryptococcus neoformans, Ustilago maydis, Rhizopus oryzae, Allomyces macrogynus, Batrachochytrium dendrobatidis, Spizellomyces punctatus, Thecamonas trahens, Dictyostelium discoideum, Polysphondylium pallidum, Entamoeba histolytica, Arabidopsis thaliana, Selaginella moellendorffii, Physcomitrella patens, Chlamydomonas* *reinhardtii, Volvox carteri* (Vcar)*, Chlorella variabilis* (Chl)*, Ostreococcus tauri* (Ostau)*, Ectocarpus siliculosus, Phaeodactylum tricornutum, Thalassiossira pseudonana, Phytophthora infestans, Toxoplasma gondii, Tetrahymena thermophila, Perkinsus marinus, Guillardia theta, Naegleria gruberi* (Ngru)*, Trypanosoma cruzi, Leishmania major, Trichomonas vaginalis, Giardia lamblia, Bigelowiella natans, Emiliania huxleyi*

- Archaea: *Nanoarchaeum equitans, Ignicoccus islandicus, Natronolimnobius baerhuensis, Halorientalis regularis, Halostagnicola kamekurae, Halalkalicoccus subterraneus, Halobiforma nitratireducens* (Halob)*, Natronobacterium gregoryi* (Natr)*, Haloplanus natans, Halovenus aranensis* (Halar)*, Halonotius pteroides* (Halo)
- Bacteria: *Actinocrispum wychmicini* (Actin)*, Kibdelosporangium aridum* (Kibd)*, Crossiella equi* (Cross)*, Lentzea xinjiangensis* (Lentz)*Nocardioides speluncae* (Noc)*, Saccharopolyspora spinosa* (Sspi)*, Synechococcus sp. PCC 7335* (Syn)*, Nostoc cycadae* (Nocyc)*, Anabaenopsis circularis* (Ancir)*, Planktothrix paucivesiculata* (Plank)*, Crinalium epipsammum* (Crinep)*, Spirosoma radiotolerans* (Spiro)*, Roseinatronobacter monicus* (Rose)

Additional BLASTp searches were conducted against a published dataset of 19 choanoflagellate transcriptomes[21], the *C. flexa* transcriptome[15] and the *Ministeria vibrans* transcriptome[70] (and courtesy of Daniel J. Richter).

Domain architectures were predicted using the CD-search tool from NCBI. For phylogenetic reconstructions, sequences were aligned using Clustal implemented in Geneious Prime (2021 version). The NOS sequence alignment was manually trimmed to be restricted to the oxygenase domain and the sGC alignment was trimmed using Gblocks with minimally stringent parameters. Phylogenetic trees were reconstructed using PhyML and BMGE implemented on http://phylogeny.lirmm.fr/phylo_cgi/index.cgi[71]. Trees were visualized using iTOL (https://itol.embl.de/)[72] and further edited in Adobe Illustrator (2021 version). Species silhouettes were added from PhyloPic (http://phylopic.org/).

**Construction of expression plasmid**

First-strand *C. flexa* cDNA (extracted as in[15]) was used as the template for cloning Cf-sGC1. Forward and reverse primers were designed against 5' and 3' ends of the target transcript (transcript name: TRINITY_DN6618_c0_g1_i1 in the published transcriptome[15]). Forward: TAAGAAGGAGATATACCATG TATGGCTTGGTGCACGAAGC; reverse: TAATGGTGATGATGGTGATG AACTATAGTCTGCTTGCCAACG. Underlined portions anneal to the sequence template. The PCR product was inserted into a pET28b vector using Gibson assembly, and the cloning product was verified by sequencing (UC Berkeley sequencing facility).

**Protein expression and purification**

pET_Cf-sGC1 was transformed into *E. coli* BL21star (DE3) cells co-expressing the chaperone GroEL/ES from the pGro7 plasmid (Takara Biosciences). After overnight incubation at 37 °C in LB Miller media supplemented with 50 µg/mL kanamycin, 20 µg/mL chloramphenicol and 500 µM iron (III) chloride, cells were subcultured 1:200 into TB media supplemented with 50 µg/mL kanamycin, 20 µg/mL chloramphenicol, 500 µM iron (III) chloride, 0.5 mg/mL L-arabinose, and 2 mg/mL glucose, grown at 37 °C. Once cell density reached $OD_{600}$ = 0.6, 1 mM 5-aminolevulinic acid was added to the culture, and culturing temperature was lowered to 18 °C. After 15 minutes of incubation, protein production was induced by addition of 500 µM isopropyl β-D-1-thiogalactopyranoside and cultures were incubated for an additional 18 hours. Cell culture was harvested by centrifuging at 4200 g for 25 min. Cells were collected, flash frozen in liquid nitrogen, and stored at -80 °C until purification.

All protein purification steps were done at 4 °C unless otherwise noted. Cells were resuspended in equal volume of buffer A (50 mM sodium phosphate, 150 mM NaCl, 5 mM imidazole, 5% glycerol, pH 8.0) supplemented with 110 mM benzamidine, 0.4 mM AEBSF, and 0.3 mg/mL DNase. Cell resuspension was lysed using an Avestin EmulsiFlex-C5 homogenizer. Cell lysate was collected and clarified by centrifugation at 32,913 g for 55 min, and the supernatant was collected and loaded onto a His60 Ni Superflow gravity column (Takara Bio). The column was washed twice, first with 10 CV buffer A, and then with 10 CV of a 9:1 mixture of buffer A and buffer B (50 mM sodium phosphate, 150 mM NaCl, 400 mM imidazole, 5% glycerol, pH 8.0). Protein was eluted with 5 CV buffer B in 1 mL fractions. Fractions with heme absorbance were pooled and concentrated using a 50kDa cutoff spin concentrator and supplemented with 5 mM DTT and 1 mM EDTA for overnight storage. Subsequently, protein was passed over a POROS HQ2 anion exchange column (Applied Biosystems). After loading, the column was washed with 5 CV of buffer C (25 mM triethanolamine, 25 mM NaCl, 5 mM DTT, pH 7.4) and developed over a gradient of 100 mM – 300 mM NaCl over 17 CV. The protein absorption spectrum was measured using a Nanodrop 2000 microvolume spectrophotometer (ThermoFisher Scientific). Subsequently, protein was aliquoted, flash frozen in liquid nitrogen, and stored at -80 °C for future use.

**Gas ligand binding of bacterial-produced Cf-sGC1**

Cf-sGC1 was handled in an argon-filled glove bag. Protein-bound heme was reduced by adding sodium dithionite (~500-fold excess over the protein conc.). Excess dithionite was removed by gel filtration of the protein into Buffer E (50 mM HEPES, 150 mM NaCl, 5% glycerol, pH 7.4) using a pre-equilibrated Zeba spin desalting column. A ferrous, ligand-free UV-vis absorption spectrum was collected on a Cary 300 UV-vis Spectrophotometer. Fe(II)-NO bound Cf-sGC1 was generated by adding the NO-releasing molecule Proli-NONOate (~10-fold excess) to the protein sample, and Fe(II)-CO bound Cf-sGC1 was generated by adding CO-sparged Buffer E to the protein sample before collecting a spectrum.

**Extinction coefficient of Cf-sGC1**

The extinction coefficient of the Soret maximum of reduced Cf-sGC1 was measured using two assays performed in tandem: heme concentration in a sample of Cf-sGC1 was determined using pyridine hemochromagen assay, and the heme Soret absorption of the protein sample was measured using UV-vis spectroscopy as described above. The pyridine hemochromagen assay was carried out following a reported protocol.[73] Briefly, a reduced protein sample with a known Soret band absorbance was diluted 5-fold in Buffer E, and then further diluted 2-fold in Solution I (0.2 M NaOH, 40% pyridine, 500 µM potassium ferricyanide) to yield the oxidized pyridine hemochromagen. An aliquot (10 µL) of Solution III (0.5 M sodium dithionite, 0.5 mM NaOH) was then added to the oxidized pyridine hemochromagen sample to yield the reduced pyridine hemochromagen. The UV-vis absorption spectrum of the reduced pyridine hemochromagen was measured on a Cary 300 UV-vis spectrophotometer. Absorption at 557 nm ($\varepsilon = 34{,}700\ mM^{-1}cm^{-1}$) was used to calculate the heme concentration in the pre-dilution sample, and the extinction coefficient of the heme cofactor of Cf-sGC1 was calculated by dividing the reduced heme absorption by the heme concentration.

**Activity assays and quantification**

Specific activity for Cf-sGC1 was measured by quantifying the amount of cGMP produced in duplicate end-point activity assays, done in biological triplicate. Cf-sGC1 from previously frozen aliquots was thawed and reduced in an anaerobic chamber as described above. The reduced protein was used without further treatment for the basal (unliganded) activity assay. A UV-vis spectrum for the unliganded sample was obtained using a Nanodrop 2000 microvolume spectrophotometer, and the Soret maximum was used to quantify heme-bound protein concentration ($\varepsilon_{428}$ = 101,000 $M^{-1}cm^{-1}$). Only protein with a Soret:280 ratio > 1 was used for activity assays. To the remaining reduced protein, proli-NONOate was added to a final concentration of 400 μM by the addition of 1 μL of a stock solution of 10 mM proli-NONOate in 10 mM NaOH, and the protein sample was incubated at 4 °C for 5 min to yield the xsNO sample. The xsNO-bound UV-vis spectrum was then collected. The protein concentration of the xsNO sample was assumed to be the same as the unliganded sample. The xsNO sample was then buffer exchanged by gel filtration using a Zeba spin column into buffer E to yield the 1-NO sample, and the UV-vis spectrum was collected. The concentration of the 1-NO sample was calculated based on the NO-bound Soret absorbance at 399 nm compared to that of the xsNO sample. Activity assays were carried out at 25 °C in Buffer E supplemented with 5 mM DTT and 3 mM $MgCl_2$ with Cf-sGC1 concentration at 40 nM. To obtain the xsNO state, 70 μM proli-NONOate was added. The reaction was initiated by addition of 1.5 mM GTP, and timepoints were quenched by diluting the reaction mixture 1:4 to a solution of 125 mM zinc acetate, and pH adjusted by adding equal volume of 125 mM sodium carbonate. Assay samples were stored at -80 °C until analyzed. Quenched assay samples were thawed at room temperature, centrifuged at 21,130 x g at 4 °C to remove zinc precipitate. Supernatant was collected and diluted 250-fold for the analysis. cGMP content of each assay sample was quantified in duplicate using an enzyme-linked immunosorbent assay (Enzo Life Sciences) following the manufacturer's protocol. Initial rate of the reaction was calculated using the linear phase of the time course, where <10% of substrate has been depleted.
